# Ancient and current association of the *Trichinella* complex with negative-sense and double-stranded RNA viruses

**DOI:** 10.64898/2026.07.06.736682

**Authors:** Gregory Karadjan, Clémence Garcia-Marin, Aurélie Heckmann, Véronique Beven, Pierrick Lucas, Yannick Blanchard, Nolwenn M Dheilly

## Abstract

The earliest records of Trichinellosis might be found in bibles that recommend against pork consumption due to the spread of a disease that resembles what is now identified as Trichinellosis. Parasitic nematodes (round worms) of the genus *Trichinella* form a complex of at least 13 species with a broad geographic range. Herein, through data mining of transcriptomic data, and re-sequencing of the transcriptome of representative isolates, we demonstrate the presence of viruses within 10 recognized *Trichinella* species. We provide genome sequences of 4 viral species of negative sense RNA viruses that belong the Family *Lispiviridae*, Order *Mononegavirales* and of 8 novel viruses of double-stranded RNA viruses that belong to a novel sub-order within the order *Ghabrivirales*. The integration of viral genome fragments within encapsulated *Trichinella* genomes demonstrate that these parasite-virus associations are ancient. Overall, viruses show co-diversification with their parasitic hosts. Yet the phylogenetic position of viruses revealed past host jump from an ancestral encapsulated *Trichinella* species to the ancestral *T. pseudospiralis*, and challenges previous dogma on the phylogeny and biogeography of *Trichinella* species in North America.

**Importance:** We are witnessing the emergence of a novel field of research focusing on viruses of parasites, and their role in parasite pathogenicity. Herein we describe the characterization of viruses within 10 recognized species of *Trichinella*. The viruses were distantly related to known viruses which led us to propose the creation of a novel genera, and a novel sub-order. We demonstrated the integration of viral genome fragments within the genome of *Trichinella* species and an on-going process of birth and death of viral integration. Finally, we showed that virus phylogenetic analyses can provide fine-tuned information of their parasitic host evolutionary history. This study demonstrates the ancient and close association between *Trichinella* and viruses, and strong species specificity. Further studies are now needed to determine whether the newly discovered viruses could be targeted for the development of novel diagnostic approaches.

## Introduction

Nematodes constitute an abundant and diverse group, with about 25000 described species, that can be found in literally any ecosystem (1). Within the clade Dorylaimia, and the order Trichocephalida, *Trichinella* is an ancient genus of parasitic nematodes specific to terrestrial carnivores with a broad geographic range and broad host species ranges. Currently, ten species and three genotypes of the genus *Trichinella* are recognized (2, 3). They are grouped in two clades that present both morphological and genotypic differences: the non-encapsulated clade that infect mammals, reptiles and birds and the encapsulated clades that infect only mammals. The encapsulated clade includes *T. spiralis* (T1); *T. nativa* (T2); *T. britovi* (T3); *T. murrelli* (T5); *T. nelsoni* (T7); *T. patagoniensis* (T12); *T. chanchalensis* (T13); and *Trichinella* genotypes T6, T8 and T9; and the non-encapsulated clade includes *T. pseudospiralis* (T4); *T. papuae* (T10); and *T. zimbabwensis* (T11), where T# denotes the genotype designation.

*Trichinella* nematodes have an auto-heteroxenous life-cycle, where all stages develop in one host, and transmission between hosts depends on predation and scavenging, sometime on the same host species. Upon consumption of infected meat containing cysts, *Trichinella* muscle larvae (ML) invade the small intestine where they develop into adult worms. Upon mating with males, females release larvae that migrate to striated muscle and encyst. *Trichinella* spp. tend to show population genetic structuring among continents but low infra-population genetic variation and low intra-specific diversity, probably due to long-distance dispersal of carnivorous hosts that promotes gene-flow within continents, and clumped transmission of siblings that promote inbreeding (4–7). Yet, natural hybrids can occur between species resulting in a discernible, but limited introgression (8, 9).

*Trichinella* also infect humans and zoonotic outbreaks of Trichinellosis occur worldwide, mostly in regions and countries where raw or undercooked meat are traditionally consumed. Trichinellosis usually begins with diarrhea, followed by fever, facial swelling, myalgia and asthenia, and can lead to fatal neurological and cardiac complications (3). The overall incidence worldwide is of about 10,000 cases / year with a mortality rate of 0.2%.

The discovery that viruses are ubiquitous in helminths and that parasite-associated viruses can contribute to associated diseases is driving an emerging paradigm (10–13) in the field. Parasitized individuals may indeed be co-infected by the parasite and the viruses carried by the parasite. Studies have demonstrated that viruses of parasites can influence the outcome of the parasitic infection by modulating parasite pathogenicity (14), parasite behavior (15), host inflammatory response (16), and host behavior (17).

While NGS-based metatranscriptomic analyses have allowed the discovery of viruses in diverse plant-parasitic nematodes (18–24), and contributed to an advanced understanding of viruses impact on the free-living nematode of the genus *Caenorhabditis* (25–27), there has been only three reports of virus discovery from animal parasitic-nematodes (28–30). Shi et al. (29) characterized the virome of a nematode of the genus *Ascaris* (order Spirurida) obtained from the gut content of pigs, a nematode of mosquitoes tentatively classified in the genus *Romanomernis* (order Mermithid), and unknown nematodes of mice, snakes and birds that belong to the class *Secernentea* (order Strongylida). A single study focused on viruses of nematodes from the order Trichocephalida: In *Capillara hepatica*, Williams et al. (28) found two viruses that belong to an unknown taxa of the order *Bunyavirales* and to the genus *Arlivirus*, family *Lispiviridae*, order *Mononegavirales*. Quek et al. (30) mostly described viruses associated with filarial nematodes *Brugia malayi* and *Onchocerca volvulus* (order Spirurida) and showed that the host develops a humoral response to filarial nematode-associated viruses. Moreover, they conducted a screen of transcriptomics data to identify viral contigs within a broad range of nematodes, including *Trichinella* sp., and identifying viral sequences belonging to the *Totiviridae* (order *Ghabrivirales*) and *Lispiviridae* (order *Mononegavirales*) families.

Herein, we used a combination of data mining and metatranscriptomic analyses to characterize viruses associated with 12 of the 13 recognized species/genotypes that constitute the *Trichinella* genus. We uncovered the complete or near-complete genomes of negative sense RNA viruses that belong to the Family *Lispiviridae*, Order *Mononegavirales* and double-stranded RNA viruses that belong to a non-recognized taxon within the order *Ghabrivirales*. Screening *Trichinella* genomes identified endogenous viral elements (EVE) that yield further insight into these ancestral *Trichinella*-virus interactions. Phylogenetic relatedness of viruses of *Trichinella* was interpreted in light of current state of knowledge on the evolutionary history of the *Trichinella* genus.

## Material and Method

### Discovery of viruses in Trichinella Transcriptomic data

To investigate the presence of viruses in *Trichinella* spp., we downloaded from the Transcriptome Shotgun Assembly (TSA) sequence database all 15 assembled transcriptomes available, representing 12 currently recognized *Trichinella* taxa, including four distinct isolates of *T. pseudospiralis* described in Korhonen et al. (2). Specific *Trichinella* isolates stored in the International Trichinella reference center (ITRC) at the Istituto Superiore di Sanità are referenced under ISS numbers. The 15 transcriptomes included *T. spiralis* (ISS3), *T. nativa* (ISS10), *T. britovi* (ISS120), *T. murrelli* (ISS417), *T. nelsoni* (ISS37), *T. patagoniensis* (ISS2496) as well as *Trichinella* genotypes T6 (ISS34), T8 (ISS272) and T9 (ISS409), *T. papuae* (ISS1980), *T. zimbabwensis* (ISS1029) and four distinct geographic isolates of *T. pseudospiralis* (ISS13, ISS588, ISS176, and ISS141) (Supp table 1). Upon identifying partial to near-complete viral genome sequences within the assembled transcriptomes, corresponding Sequence Read Archive (SRA) files were downloaded and assembled for virus discovery purpose as described below.

**Table 1:** Endogenous Viral Elements of Totiliviruses were found within long genome scaffolds.

| Species | strain | Accession | scaffold | length (bp) | Coordinates | hit length (bp) | Name |
| --- | --- | --- | --- | --- | --- | --- | --- |
| *T. britovi* | ISS120 | JYDI01000062.1 | scaffold62s | 195695 | c61913-61408 | 509 | CP7 |
| *T. murrelli* | ISS417 | JYDJ01000189.1 | scaffold189s | 87692 | 32493-33002 | 509 | CP7 |
|  | ISS35 | LKDY01000594.1 | unitig_1216 | 1425034 | 833434-833943 | 509 | CP7 |
| *T. sp. T6* | ISS34 | JYDK01000012.1 | scaffold12s | 399341 | 352514-353023 | 509 | CP7 |
| *T. sp. T9* | ISS409 | JYDN01000063.1 | scaffold63s | 211722 | 194989-195498 | 509 | CP7 |
| *T. patagoniensis* | ISS2496 | JYDQ01000167.1 | scaffold167s | 97436 | 1923-2468 | 545 | RdRp3 |
| *T. nativa* | ISS10 | JYDW01000003.1 | scaffold3s | 395314 | c47674-47165 | 509 | CP7 |
|  | ISS11 | JYDW01000017.1 | scaffold17s | 311569 | c264335-263700 | 635 | RdRp2 |

### Sample collection and RNA extraction

Seventeen isolates of first-stage muscle larvae (ML) representing 12 of the recognized species and genotypes of *Trichinella* were produced at the ANSES Parasite National reference Laboratory, Maisons-Alfort, France. The seventeen isolates included *T. spiralis* (ISS004 and ISS7216), *T. nativa* (ISS10, ISS42 and ISS70), *T. britovi* (ISS1721 and ISS 7732), *T. murrelli* (ISS417), *T. nelsoni* (ISS37), *T. patagoniensis* (ISS1826) as well as *Trichinella* genotypes T6 (ISS34), T8 (ISS124) and T9 (ISS408), *T. papuae* (ISS572), *T. zimbabwensis* (ISS1029) and *T. pseudospiralis* (ISS13, ISS470) (Supp table 1).

*Trichinella* spp. were maintained by several passages in 5-week-old female OF-1 mice (Charles River laboratories, France). The mice were maintained under specific pathogen-free conditions in individual closed cages on a ventilated rack. Infected mice carcasses were artificially digested to obtain *Trichinella* spp. L1 as previously described (31).

RNA extractions were performed using the RNAqueous – Micro Kit® (Ambion, France) following the manufacturer’s recommendations. Firstly, 1000 L1M and 50 µL of lysis buffer were placed in VK05 tubes (Ozyme, France) and crushed in a Bead Beater (Precellys, France) using three crushing cycles at 6500 rpm for 15 seconds with breaks of 10 s. The tubes were then rinsed with an additional 50 µL of lysis buffer. Fifty microliters of absolute ethanol was then added to the crushed mixture and quickly vortexed. Of the resulting solution, 150 µL was placed in the extraction column and centrifuged for 30 s at 15000 g. The column was washed once in Wash Solution 1 then twice in Wash Solution 2/3 with centrifugation for 30 s at 15000 g at each step. Then the columns were centrifuged for 1 min at 15000 g to further dry the membrane. Finally, RNA was eluted by adding 10 µL of RNase-free water, 1 minute incubation period at room temperature and centrifuging for 10 s at 15000g. This was done twice.

DNase treatment was then performed according to the manufacturer’s recommendations (RNAaqueous-Micro kit, Ambio, France). An initial mixture of 2µl ‘10X DNase I Buffer’ and 1µl ‘DNase I’ was made and added to the RNA sample. The tubes were incubated at 37°C for 20 minutes. Then 2µl of ‘DNase Inactivation Reagent’ was added and incubated for 2 minutes at room temperature. The tubes were centrifuged at 15000g for 1 minute and 30 seconds. Finally, the supernatant was collected.

### Transcriptome sequencing

The cDNA libraries were prepared with the Illumina Stranded Total RNA Prep, Ligation with Ribo-Zero Plus Kit (Illumina, Évry-Courcouronnes, France) according to the supplier’s instructions. The cDNA libraries were sequenced using a NovaSeq Sequencer (Illumina) with the NovaSeq 6000 SP reagent kit (300 cycles). Raw reads for RNA-sequencing were submitted in the NCBI BioProject database with accession PRJNA1193013

### Viral contigs discovery

All SRA data, and in-house sequenced transcriptomes were assembled in-house and used to search for viral genomic sequences as described previously (11). Briefly, sequences were trimmed for adapter removal using Fastp version 0.20.1 (32) sequence quality was verified using FastQC version 0.11.8. Sequence reads were then mapped against the corresponding *Trichinella* genome using Bowtie 2 (33). Unmapped reads were assembled with Mira (version 4.0.2) and SPAdes (version 3.10.0) *de novo* assembler (34, 35). The resulting contigs were aligned on local nt database composed of *Trichinella* genomes with megablast (version 2.10.0, options: "-perc_identity 90 -max_target_seqs 1") to remove host sequences. Contigs larger than 500 bp were initially searched for viral hits using Megablast against the NCBI nonredundant (nr) nucleotide database, with no success. Viral contigs were discovered in TSA and assembled SRA data by comparing (Blastx, E value of < 10^-10^) against a viral protein database containing sequences from the NCBI database (RefSeq release number 93, https://www.ncbi.nlm.nih.gov/refseq/). To confirm virus discovery, putative viral transcripts were translated into proteins to conduct reciprocal blastp against Genbank nonredundant (nr) protein database 244, released on June 25^th^, 2021. Reassembly efforts used a combination of Blast searches and reads alignments using Burroughs-Wheeler Aligner (BWA, version 0.7.8)(36) and visualization in Integrative Genome Viewer (IGV)(37, 38), to complete gaps between contigs and/or extend the length of partial genome sequences. The list of novel viruses identified, best blast hit, taxonomic assignment, and completeness of coding regions is provided in Supp table 2.

### Phylogenetic analyses

Predicted open reading frame (ORF) boundaries were extracted using Prokka (39). The viral predicted proteins and polyproteins were then aligned to closely related viruses to confirm completeness and provide assistance with viral genome annotation. Annotation of domains and initial supergroup assignation was extracted from comparisons against the Conserved Domain Database (CDD) as implemented by BLASTp against the nr protein database. Pairwise identity matrices were obtained using Clustal-Omega online (https://www.ebi.ac.uk).https://www.ebi.ac.uk). Viral RNA-dependent RNA polymerase (RdRp) sequences were aligned, using the E-INS-I algorithm implemented in the program MAFFT (version 7), to representative sequences of all viral families and genera ratified by the ICTV, as well as closely related viruses identified through blast searches(40). Ambiguously aligned regions were removed using TrimAl (version 1.2) (41). For each data set, the best-fit model of amino acid substitution was determined using Smart Model Selection (SMS) as implemented in PhyML (version 3.0) (42). Phylogenetic trees were then inferred using the maximum-likelihood method implemented in PhyML (version 3.0) using the best-fit model and best of Nearest-Neighbor Interchange (NNI) and Subtree Pruning and Regrafting (SPR) branch swapping (42). Support for nodes on the trees was obtained using an approximate likelihood ratio test (aLRT) with the Shimodaira-Hasegawa-like procedure. Viruses were taxonomically classified based on their phylogenetic positions, and pairwise sequence comparisons (43).

### Endogenous viral elements

Genomes of *Trichinella* (Taxid: 6333) were screened for the presence of viruses using blastn and tBlastn with either viral genome sequences or predicted proteins as queries and a cutoff value of 1e^-10^. A total of 24 Whole Genomes Sequencing databases were screened*: T. spiralis* (JYDH01, JBEUSY01), *T. britovi* (JYDI01), *T. murrelli* (JYDJ01), T. sp. T6 (JYDK01), *T. nelsoni* (JYDL01, JAOTOD01), T. sp. T9 (JYDN01), *T. patagoniensis* (JYDQ01), *T. pseudospiralis* (JYDR01, JYDU01, JYDT01, JYDS01, JYDV01, QAWF01, JBEUSZ01), *T. spiralis* (ABIR03), T. sp. T8 (JYDM01), *T. papuae* (JYDO01), *T. zimbabwensis* (JYDP01), *T. nativa* (JYDW01, LVZM01), *T. murrelli* (LKDY01), and T. sp. 17WV049-YT159. The potential EVEs were extracted from the genomic contigs and scaffolds and filtered using a reciprocal blastn against NCBI nr database. EVE identified are provided in Supp Table 3. The filtered EVEs were aligned to reference exogeneous viruses of *Trichinella* and included in Phylogenetic analyses as described above.

### Polymerase chain reactions

Based on the sequence obtained by NGS, and for each viral species, primer pairs were designed using Primer 3 Plus (44) (supp table 4). When pools of different isolates of *Trichinella* from the same species, or pools of different genotypes of *Trichinella* were prepared before NGS sequencing, the host assignment of the newly identified viruses was determined based on the PCR results and on viral sequence identities to viruses detected in other datasets as indicated in Supp Table 1.

Reverse transcription was performed using the Maxima Reverse Transcription Kit (Thermo, France) as recommended. Briefly, we used 4 µL of 5X RT Buffer, 1 µL of Oligo(dT), 1 µL of dNTP, 0.5 µL of RiboLock RNase Inhibitor, 9 µg of sample RNA sample in a total volume of 20 µL. Complementary DNA (cDNA) were obtained after 30min incubation at 50°C and 5min at 85°C. The Polymerase chain reaction was performed using Taq Phusion within a 50 µL final volume composed of 10 µL 10X buffer, 1 µL dNTP, 2.5 µL of each primer at 10µM (supp table 2), 0.5 µL of Taq DNA polymerase and 10 µL of cDNA. The PCR consisted in Taq activation for 2min at 98°C followed with 40 cycles of denaturation for 10s at 98°C, hybridization for 1 min at 60°C and elongation for 1min at 72°C and finished by an elongation during 10min at 72°C. PCR products were visualized on a 2% agarose gel. The specificity of the amplification was confirmed by sequencing of gel bands of the expected size range.

### Data availability

Viral sequences are available under GenBank accession numbers PQ673584-PQ673600 as indicated in Supp table 2.

### Ethical statements

All animal experiments were carried out in accordance with EU Directive 2010/63/EU and French legislation, namely Decree no. 2013-118 of 1 February 2013 issued by the French Ministry of Agriculture, Agrifood and Forestry. The ethical committee approved all the experiments: C2EA-16 Comité d’éthique ComEth ANSES/ENVA/UPEC, under the following approval number: saisine 12-0048, ComEth 13/11/12-4.

## Results

### Two taxa of viruses within the *Trichinella* species complex

We screened for viruses *in* a total of 25 RNA sequencing datasets representing 27 distinct isolates of all 12 currently recognized *Trichinella* taxa (Supp table 1). These included 15 isolates / 12 species described in Korhonen et al. (2) and 17 isolates / 12 species sequenced as part of this study. Sequencing data were assembled *de novo* and processed as previously (11) for viral contigs discovery. Viruses were found associated with *T. patagoniensis*, *T. nativa*, T. sp. T6, T. sp. T8, T. sp. T9, *T. murrelli*, *T. britovi* and *T. pseudospiralis*, whereas no virus was discovered within *T. spiralis*, *T. nelsoni*, *T. papuae* and *T. zimbabwensis*. Novel viruses belonging to the family *Lispiviridae* and novel viruses of the order *Ghabrivirales* were found in 3 and 8 different species of *Trichinella*, respectively. Closely related viruses were uncovered in both encapsulated and non-encapsulated *Trichinella* species suggesting common origin. In fact, within a given taxon, all discovered viruses were closely related, and displayed similar length and the same genome structure and composition (Figure 1 and 2). The broad species distribution of the newly discovered viruses prompted us to compare the viruses phylogenetic trees with *Trichinella* evolution inferred from phylogenomics and phylogeography (2). Moreover, using the newly discovered viral genomes as reference, a search for Endogenous viral elements (EVE) led to the identification of EVEs covering partial regions of the Capsid and RdRp of the ghabriviruses providing additional information on the viruses’ evolutionary history (Figure 1, Supp table 3).

**Figure 1:**
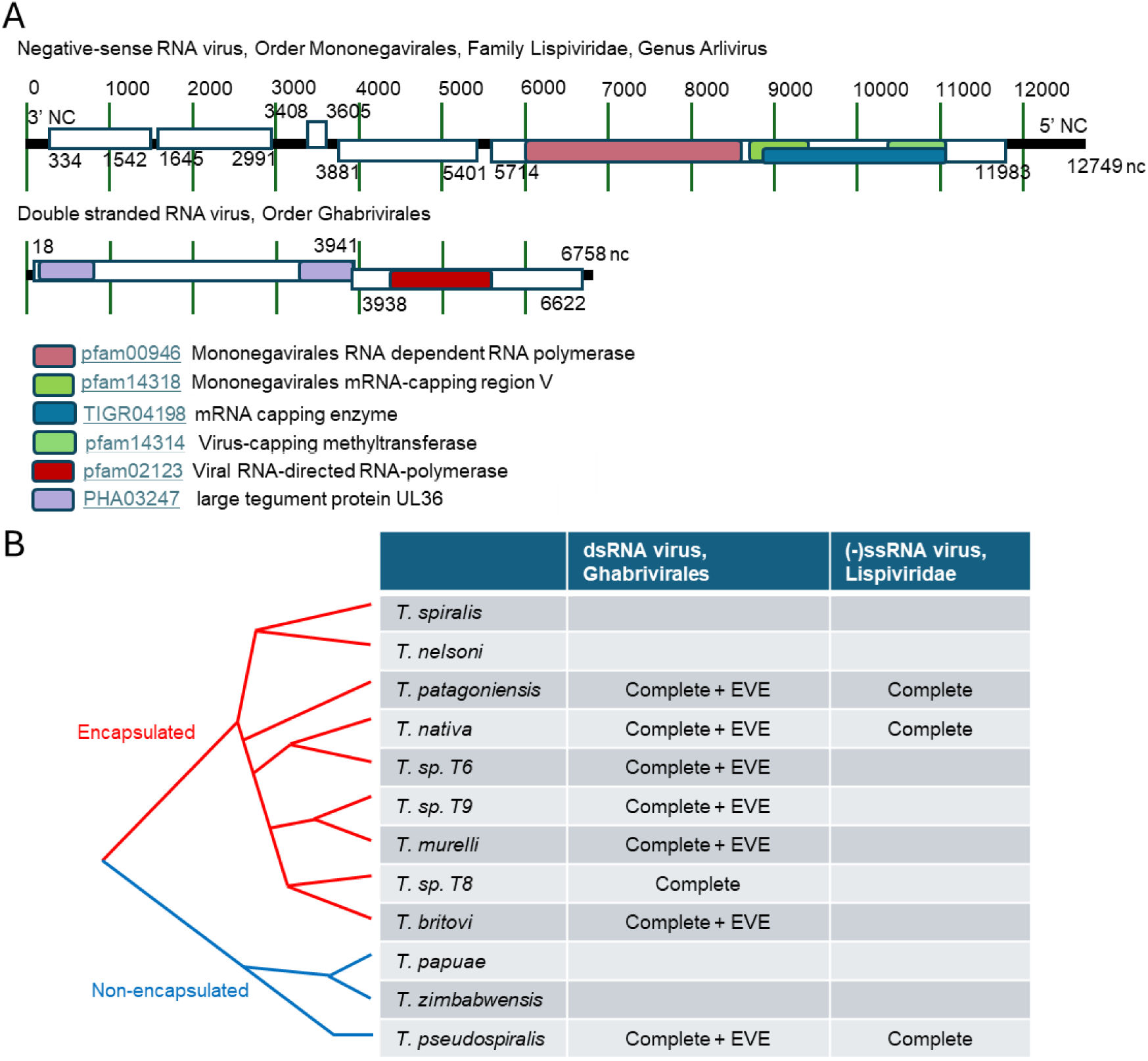
Discovery of viruses within *Trichinella*. A/ Genome composition of exemplar viruses from *T. pseudospiralis*. B/ The phylogeny of the 12 currently recognized taxa of *Trichinella* as described by Korhonen et al. 2016, and viruses identified. No exogenous nor endogenous viruses were found in *T. spiralis*, *T. nelsoni*, *T. papuae* or *T. zimbabwensis*.

**Figure 2:**
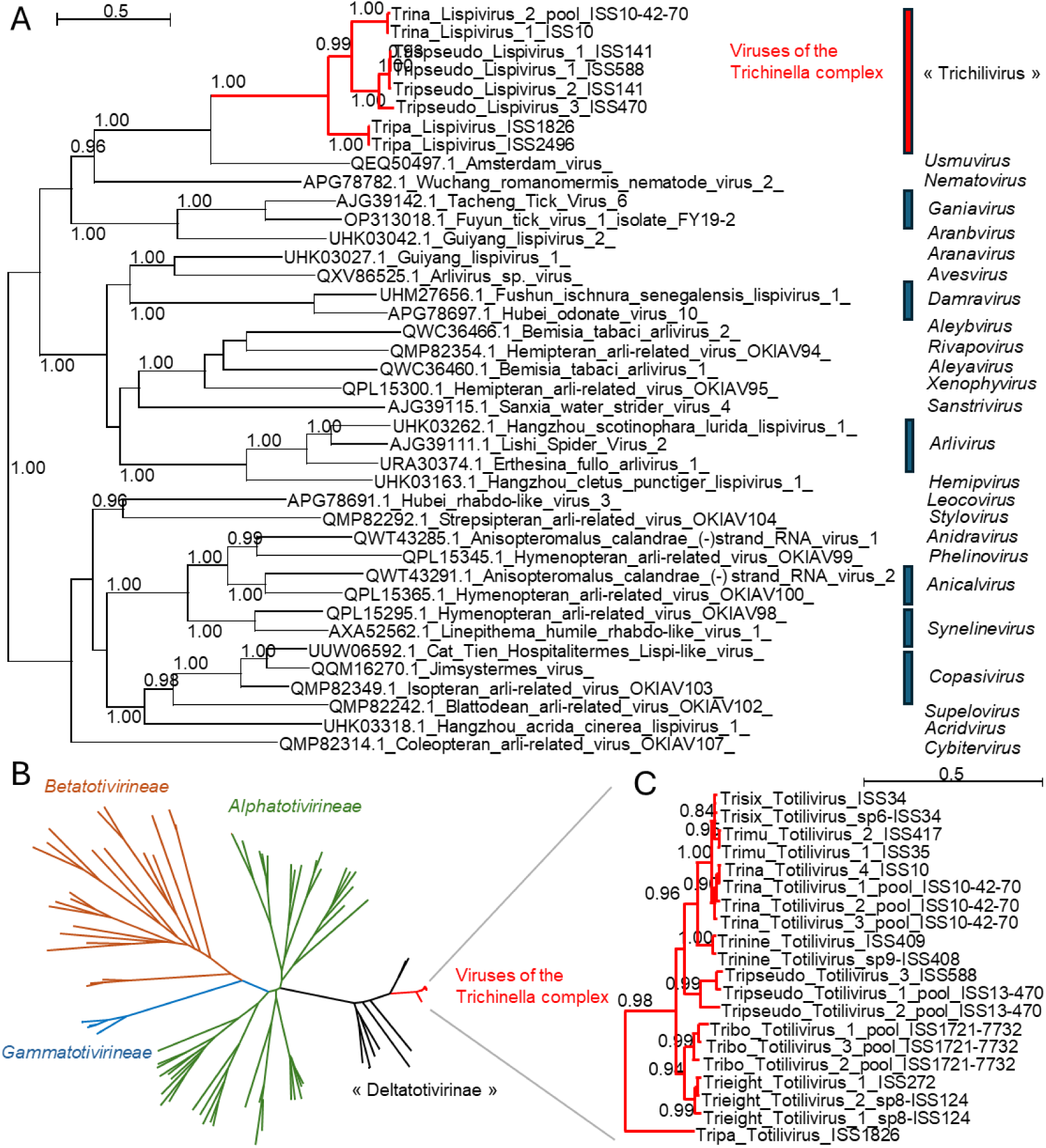
Viruses of *Trichinella* belong to two novel taxa. A/ Phylogenetic tree of the RNA-dependent RNA polymerase (RdRp) of RNA viruses of the family *Lispiviridae*, order *Mononegavirales*. Viruses of *Trichinella* constitute a novel genus tentatively named “*Trichilivirus*”. B/ The phylogenetic analysis of the RdRp of RNA viruses of the order *Ghabrivirales* reveals the existence of a novel sub-order tentatively named “*Deltatotivirineae*”. C/. Viruses of the *Trichinella* complex constitute a novel taxon within the proposed *“Deltatotivirineae*” sub-order. All trees were inferred in PhyML using the LG substitution model. Branch points indicate that results of Shimodaira-Hasgawa branch test > 0.9.

### A novel genus within the family *Lispiviridae with four species*

Within *T. nativa*, *T. patagoniensis* and *T. pseudospiralis*, we found closely related viruses with 55-56% identity with the Amsterdam virus which belongs to the genus *Usmuvirus,* family *Lispiviridae,* order *Mononegavirales* of negative-sense RNA viruses. (Supp Table 2). Phylogenetic analyses on the complete RdRp sequence confirmed that these viruses belong to the Family Lispiviridae. In fact, based on the family criteria, and with ≥50% RdRp amino acid identity, all lispiviruses of *Trichinella* constitute together a novel genus tentatively named *“Trichilivirus*” (Figure 2, supp table 5).

Both the RdRp protein phylogenetic analysis (Figure 2) and the full-length genome sequence phylogenetic analyses (not shown) showed that the Tripa Lispivirus identified in *T. patagoniensis* has a basal position to viruses of *T. pseudospiralis* and *T. nativa.* According to ICTV, lispiviruses that share ≥85% RdRp amino acid identity are considered to be the same species. Each species of *Trichinella* was found to host a single species of trichilivirus, except for the Tripseudo Lispivirus 3 of *T. pseudospiralis* ISS470 that is sufficiently divergent from both Tripseudo Lispivirus 1 (from *T. pseudospiralis* ISS 588 and ISS141) and Tripseudo Lispivirus 2 (from *T. pseudospiralis* ISS141) to constitute a distinct species.

### A new viral species, within a novel sub-order within the order *Ghabrivirales*

Within *T. nativa*, *T. patagoniensis*, *T. britovi*, *T murrelli*, *T. pseudospiralis*, T. sp T6, T. sp T8 and T. sp T9, we found closely related viruses with 55-56% identity to the unclassified Nanning Totiv tick virus (Supp Table 2) within the order *Ghabrivirales*. Phylogenetic analyses on the complete RdRp sequences revealed that viruses of *Trichinella* do not belong to any of the sub-orders recently recognized by ICTV: the *Alphatotivirineae*, the *Betatotivirineae* and the *Gammatotivirineae*. Viruses of *Trichinella* cluster on the tip of a separate branch that includes only unclassified viruses, which prompted us to propose a fourth sub-order, tentatively named *“Deltatotivirineae*” (Figure2). All viruses within this sub-order have an undivided typical dsRNA genome with two separate Open Reading Frames (ORFs) that encode for the CP and RdRp, respectively. Further supporting the novel sub-order classification, these viruses show a high-level of RdRp amino-acid sequence identity (supp table 6) and show distinct consensus RdRp motif sequence from other suborders (Figure 3). According to ICTV, ghabriviruses that share ≥70% RdRp amino acid identity are considered to be the same species, and those that share ≥31% RdRp amino acid identity belong to the same genus. Following this classification criteria, all novel ghabriviruses of *Trichinella* belong to the same species tentatively named Totilivirus. Together with Nanning totiv tick virus 4 and Hubei toti-like virus 6, the three viruses constitute a novel genus. All other unique species within the “*Deltatotivirineae*” appear to be sole members of their genera, but with more than 21% RdRp amino acid identity, they can all be considered to belong to the same family.

**Figure 3:**
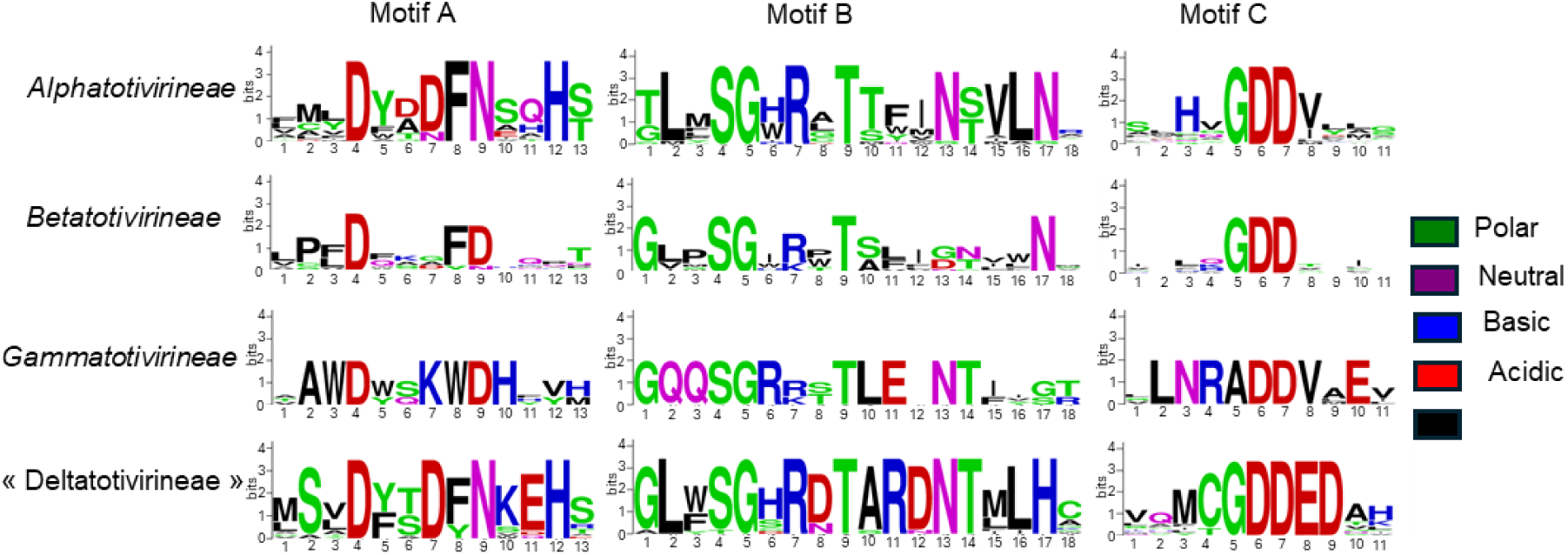
Consensus amino acid sequences of the RdRp motifs of the proposed “Deltatotivirinaeae” suborder compared to recognized sub-orders within the Order *Ghabrivirales.* The motifs A, B, C were visualized by WebLogo (v3.7.12 REF).

Full*-*length viral genome alignments and phylogenetic analyses were conducted in order to investigate the evolutionary history of the Totilivirus within the *Trichinella* species complex (Figure 4). Overall, the virus evolution agrees with the phylogenetic relationship between *Trichinella* species further suggesting co-diversification, with two major exceptions. First, the virus of *T. patagoniensis* has a basal position to viruses of non-encapsulated *T. pseudospiralis* and to viruses of encapsulated *Trichinella* species. Secondly, viruses of *T. murrelli* are more closely related to viruses of T. sp T6 whereas phylogenomics of the *Trichinella* complex shows pairs of sister species (*T. nativa* + T. sp T6, *T. murrelli* + T. sp T9 and *T. britovi* + T. sp T8).

**Figure 4:**
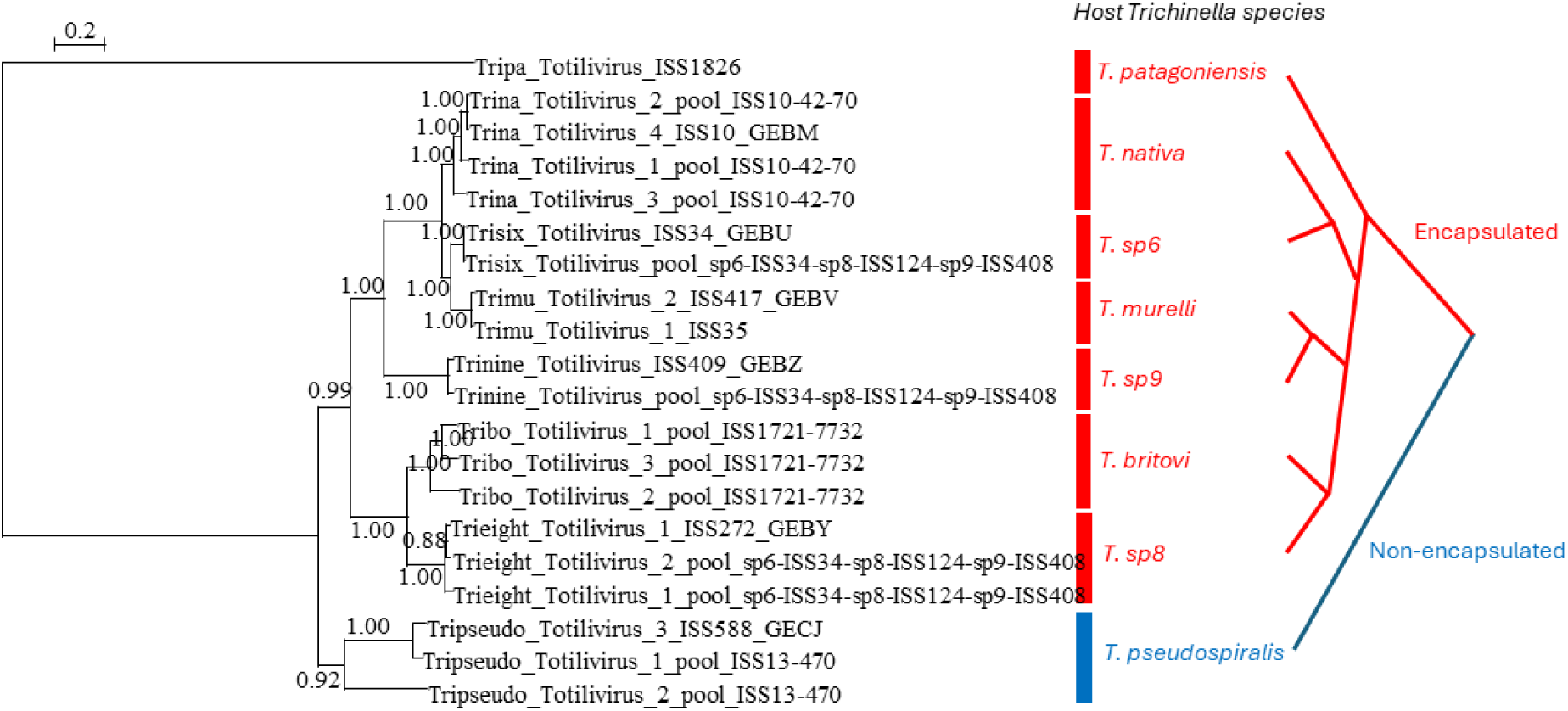
Comparison of the phylogenetic relationship among variants of Totiliviruses identified in different *Trichinella* species, and their host phylogenetic relationship as described in Korhonen *et al*. (2016). The figure provides a phylogenetic tree of the full-length genomes of Totiliviruses. The tree was inferred in PhyML using the LG substitution model. Branch points indicate that results of Shimodaira-Hasgawa branch test > 0.9.

### Birth and death of endogeneous viral elements of totiliviruses

Endogenous Viral Elements of Totiliviruses were found within the genomes of *T. patagoniensis, T. nativa*, T. sp. T6, T. sp. T9*, T. murrelli*, *T. britovi* and *T. pseudospiralis*. The viral hits covered either a partial capsid protein (up to seven homologous EVE), or a partial RdRp (up to 3 homologous EVEs) (table 1). All identified EVEs but CP7-EVE appear to have been recently endogenized given their very close phylogenetic relationship with identified exogeneous Totiliviruses associated with the same *Trichinella* species (Supp fig 1). Most contigs or scaffolds were too short to further characterize the EVEs beyond phylogenetic positioning (Supp table 3 and Supp fig 1). Yet, we could localize the RdRp2-EVE on scaffold17s of *T. nativa* (JYDW01000017.1, table 1), in a region flanked by PGBD2 (mRNA Piggy-Bac transposable element derived protein 2) and smarca1 genes, and the RdRp3-EVE on scaffold167s of *T. patagoniensis* (JYDQ1000167.1, table 1) in a region flanked by hypothetical proteins and Tigd2 (mRNA Tigger transposable element derived protein 2). Blastn of the genomic regions against other *Trichinella* genomes confirmed the absence of orthologous EVEs, further demonstrating that the RdRp2-EVE and RdRp3-EVE have been recently integrated within the genome of *T. patagoniensis* and *T. nativa* respectively.

The CP7-EVE was most closely related to the virus of *T. patagoniensis* (Supp figure 1) and was found in several encapsulated *Trichinella* species suggesting that it was endogenized in a shared common ancestor (table 1). We confirmed orthology of the CP7-EVE in *T. nativa* (scaffold 3s, JYDW01000003.1), *T. britovi* (scaffold62s, JYDI01000062.1), T. sp. T6 (scaffold12s, JYDK01000012.1), *T. murrelli* (scaffold189s, JYDJ01000189.1), T. sp. T9 (scaffold63s, JYDN01000063.1): the CP-EVEs were all flanked by iff-2 (mRNA eukaryotic translation initiation factor 5A-2) and rps-26 (mRNA 40S ribosomal protein S26) genes. Within T. sp. T8 the EVE initially appeared absent. Yet, after finding the orthologous scaffold enclosing iff-2 and rps-26 (scaffold89s, JYDM01000089.1), the alignment was indicative of a past endogenization of CP7-EVE, followed with a deletion: a 34nt long fragment of CP7-EVE remained, while a fragment of the flanking region was deleted (figure 5). In contrast, we confirmed CP7-EVE absence of orthologous genomic regions in the genomes of *T. spiralis* (tig00000734, JBEUSY010000170.1), *T. nelsoni* (scaffold25s, JYDL01000025.1), and *T. patagoniensis* (scaffold1s, JYDQ01000001.1). This analysis allows us to infer that the genetic fixation of CP7-EVE likely occurred after the divergence of *T. patagoniensis*, and before the encapsulated *Trichinella* taxa expanded.

**Figure 5:**
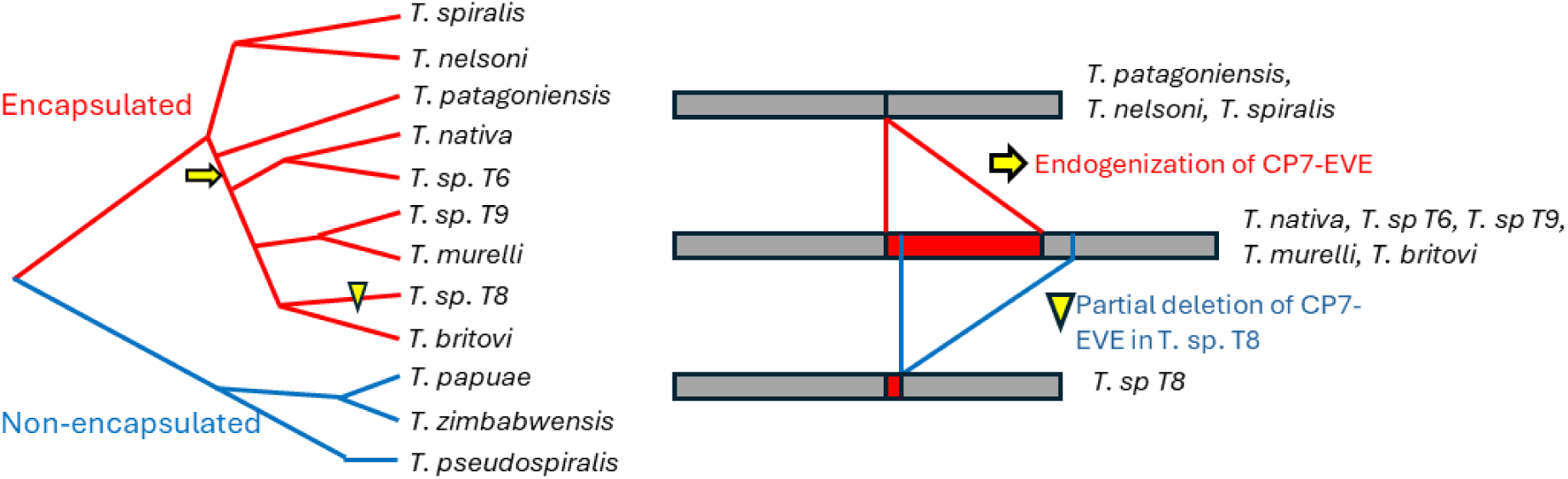
proposed history of endogenization and deletion of EVE-CP7 in encapsulated *Trichinella* species.

## Discussion

The study of viruses of parasites is a very novel and expanding field of research, with broad applications. Herein, we combined metatranscriptomics analysis and paleovirology to characterize current and past viral infections in the *Trichinella* complex. Positioning of the novel viruses within the known viral diversity led to the characterization of several novel viral taxa, including a new sub-order within the order *Ghabrivirales*, and a new genus within the family *Lispiviridae*. Diverse dsRNA viruses of the order *Ghabrivirales* have been found associated with parasitic species such as the cestode *Schistocephalus solidus* (45) and a diversity of trematodes (11), and protozoan parasites such as *Leishmania*, *Giardia* and *Trichomonas*. dsRNA of parasites were found to contribute to host inflammation and susceptibility to parasite infection (14, 16, 46), which justify further studies on the role of Totiliviruses in *Trichinella* pathogenicity. Lispiviruses have currently been found mainly associated with arthropods and nematodes and their transmission to parasitized hosts have not been assessed.

The viruses phylogenetic diversity within the *Trichinella* complex were interpreted in light of current knowledge of the evolutionary history of *Trichinella*, as reconstructed via phylogenomics analyses by Korhonen et al. (2). We found closely related viruses in both encapsulated (*T. patagoniensis*, *T. nativa*, T. sp. T6, *T. murelli*, T. sp. T9, *T. britovi* and T. sp. T8) and non-encapsulated (*T. pseudospiralis*) *Trichinella* species indicating shared ancestry. Neither exogenous viruses nor EVE were found in encapsulated ancestral species *T. spiralis* and *T. nelsoni*, nor in non-encapsulated *T. papuae* and *T. zimbabwensis*. While we cannot exclude the possibility of biased sampling, one explanation would be that both the Trichilivirus and Totilivirus emerged in Paleartic between 7 and 5 mya, after the first biotic expansion that separated *T. pseudospiralis* from the lineage with *T. papuae* + *T. zimbabwensis*, and resulted in the expansion of *T. nelsoni* + *T. spiralis* lineage. This hypothesis is supported by the close phylogenetic relatedness of CP7-EVE with the Totilivirus of *T. patagoniensis*, and the absence of EVE within these ancestral *Trichinella* species.

For both viral taxa, phylogenetic relatedness agrees broadly with the state of knowledge in *Trichinella* evolutionary history, with a clear co-diversification of *Totili*viruses with their parasitic hosts. The observed co-diversification likely results from the strong geographic and ecologic isolation of *Trichinella* species. Yet, we identified two major discrepancies that provide valuable information on *Trichinella* parasite and virus evolutionary history.

First, our analyses showed that both the Trichiliviruses and Totiliviruses, viruses of *T. patagoniensis* have a basal position to viruses of both encapsulated and non-encapsulated *Trichinella* species. The most parsimonious explanation is probably a viral emergence in an ancestral encapsulated *Trichinella* followed with viruses’ host jump around 5 Mya ago, after the divergence of *T. patagoniensis*, and before the encapsulated *Trichinella* taxa expanded and diversified. Virus host jump would require that different *Trichinella* species share the same geographic range and infect the same host species. Hybridization as observed in experimental and field conditions (8, 47, 48) may have allowed this host jump. Interestingly, the proposed host jump match well with the biogeography of *Trichinella.* Around 5-7 mya, the shared ancestor of encapsulated *Trichinella* and the shared ancestor of *T. pseudospiralis* isolates would have used wild boars and mountain lions as hosts and were localized around the palearctic (2, 48, 49).

The second discrepancy between virus phylogenetic analyses and *Trichinella* evolutionary history concerns the observed relatedness of the Totiliviruses of *T. murrelli* and T. sp. T6 (also reported by Quek et al. 2024 from partial genome sequences) whereas phylogenomics of the *Trichinella* complex revealed pairs of sister species (*T. nativa* + T. sp T6, *T. murelli* + T. sp T9). Yet, other studies led to divergent conclusions regarding the history of encapsulated *Trichinella* speciation events (48, 49) and the succession of events that led to T. sp. T9, *T. murrelli*, *T. nativa* and T. sp. T6 divergence remain unresolved. We suggest that the evolutionary history of the totiliviruses may allow a better fine-tuning of the evolutionary history of encapsulated *Trichinella* species. The virus evolutionary history agrees with the initial divergence of *T. britovi* and T. sp. T8 on the African continent (3.2-1.7 mya), and the secondary expansion of Nearctic and Eurasian obligate carnivory *Trichinella* species. It also suggests an initial divergence of T. sp. T9 from a common ancestor that migrated to Japan, whereas the common ancestor of *T. nativa*, T. sp. T6 and *T. murrelli* crossed the Bering land bridge. Phylogenetic distance between totiliviruses would then agree with a secondary geographic isolation of *T. nativa*, T. sp. T6 and *T. murrelli* along a gradient from North to South with *T. nativa* representing a persistent high latitude population from which T. sp. T6 and *T. murrelli* diverged when moving south.

To conclude, this study not only provide a thorough characterization and classification of the viruses of *Trichinella*, but we also provide evidence of ancient and persistent association between viruses and *Trichinella* species and demonstrate that parasite-associated viruses phylogenetic analysis can provide information on the parasite evolutionary history. Further studies are needed to characterize virus replication kinetics and determine the extent to which viruses of *Trichinella* contribute to the host-parasite interaction. The recent demonstration that parasitized hosts can produce antibodies directed against viruses of several helminth species (30, 50) suggests that viruses could contribute to parasite virulence or modulate host susceptibility to infection.

## Acknowledgments

This project is part of the Parasite Microbiome project and was funded by the French Agency for Food, Environmental and occupational Health Safety project PARAVIR to NMD and GK.

## Notes

### Competing Interest Statement

The authors have declared no competing interest.

