## supplementary figure 1 for "Ancient and current association of the *Trichinella* complex with negative-sense and double-stranded RNA viruses"

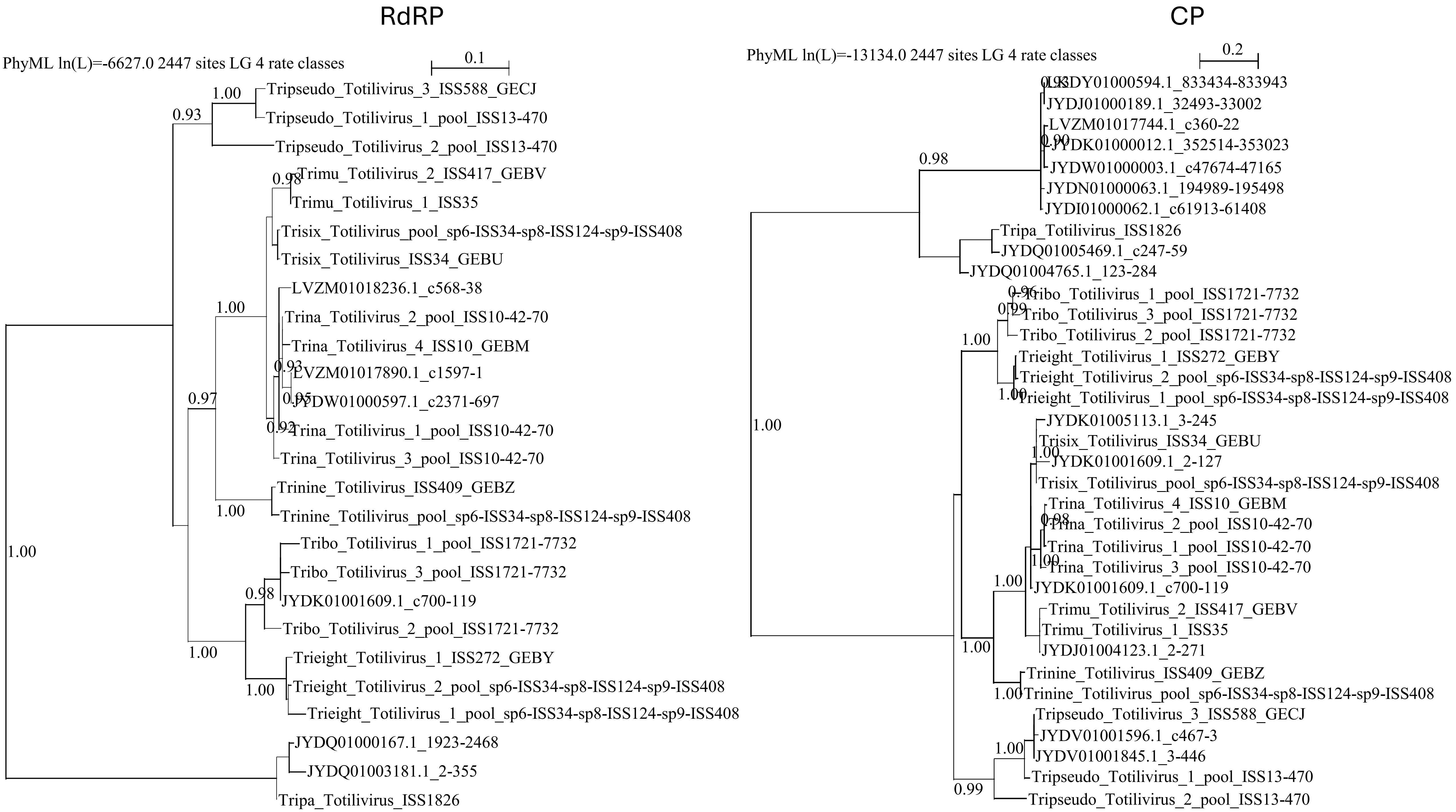


Supp Fig 1: Comparison of the phylogenetic relationship among proteins encoded by EVE and Totiliviruses identified in different *Trichinella* species. Phylogenetic tree of the RNA-directed RNA polymerase (RdRP) and Capsid (CP) proteins. The trees were inferred in PhyML using the LG substitution model. Branch points indicate that results of Shimodaira-Hasgawa branch test > 0.9.
